# A Platform Incorporating Trimeric Antigens into Self-Assembling Nanoparticles Reveals SARS-CoV-2-Spike Nanoparticles to Elicit Substantially Higher Neutralizing Responses than Spike Alone

**DOI:** 10.1101/2020.06.11.147496

**Authors:** Baoshan Zhang, Cara W. Chao, Yaroslav Tsybovsky, Olubukola M. Abiona, Geoffrey B. Hutchinson, Juan I. Moliva, Adam S. Olia, Amarendra Pegu, Emily Phung, Guillaume Stewart-Jones, Raffaello Verardi, Lingshu Wang, Shuishu Wang, Anne Werner, Eun Sung Yang, Christina Yap, Tongqing Zhou, John R. Mascola, Nancy J. Sullivan, Barney S. Graham, Kizzmekia S. Corbett, Peter D. Kwong

**Affiliations:** Vaccine Research Center, National Institute of Allergy and Infectious Diseases, National Institutes of Health, Bethesda, Maryland, USA; Electron Microscopy Laboratory, Cancer Research Technology Program, Leidos Biomedical Research Inc., Frederick National Laboratory for Cancer Research, Frederick, Maryland, USA; Institute for Biomedical Sciences, George Washington University, Washington, DC, USA

## Abstract

Antigens displayed on self-assembling nanoparticles can stimulate strong immune responses and have been playing an increasingly prominent role in structure-based vaccines. However, the development of such immunogens is often complicated by inefficiencies in their production. To alleviate this issue, we developed a plug-and-play platform using the spontaneous isopeptide-bond formation of the SpyTag:SpyCatcher system to display trimeric antigens on self-assembling nanoparticles, including the 60-subunit *Aquifex aeolicus* lumazine synthase (LuS) and the 24-subunit *Helicobacter pylori* ferritin. LuS and ferritin coupled to SpyTag expressed well in a mammalian expression system when an *N-*linked glycan was added to the nanoparticle surface. The respiratory syncytial virus fusion (F) glycoprotein trimer – stabilized in the prefusion conformation and fused with SpyCatcher – could be efficiently conjugated to LuS-SpyTag or ferritin-SpyTag, enabling multivalent display of F trimers with prefusion antigenicity. Similarly, F-glycoprotein trimers from human parainfluenza virus-type 3 and spike-glycoprotein trimers from SARS-CoV-2 could be displayed on LuS nanoparticles with decent yield and antigenicity. Notably, murine vaccination with the SARS-CoV-2 spike-LuS nanoparticles elicited ~25-fold higher neutralizing responses, weight-per-weight relative to spike alone. The versatile platform described here thus allows for multivalent plug-and-play presentation on self-assembling nanoparticles of trimeric viral antigens, with SARS-CoV-2 spike-LuS nanoparticles inducing particularly potent neutralizing responses.

## Introduction

Self-assembling nanoparticles are playing an increasingly prevalent role in vaccine development as vaccine vehicles and immunomodulators. The appeal of nanoparticle immunogens lies in their inherent multivalent display of antigens, which is known to elicit robust B cell responses (reviewed in ^1^). There have been numerous efforts to genetically fuse viral immunogens to nanoparticles within prokaryotic and eukaryotic systems, utilizing direct genetic fusion of antigenic molecules with self-assembling nanoparticle monomers^2–4^, chemical conjugation^5,6^, and a spontaneous intramolecular isopeptide bond formation with the SpyTag:SpyCatcher system^7,8^, and some of these nanoparticles are now entering clinical trials^2,9,10^.

Another important factor to consider in viral immunogen design is glycosylation. Viral pathogens are often heavily glycosylated, often as a means to evade the human immune system. Moreover, many viral antigens require glycosylation to be stably expressed and correctly folded. Although several studies have described plug-and-play nanoparticle systems^11–16^, many use prokaryotic expression systems, which are not suitable to produce correctly glycosylated antigens. Furthermore, *N*-glycans can be manipulated in immunogen design to selectively occlude unwanted epitopes as well as to improve the solubility and stability of immunogens^17–19^. Another factor to consider is that metastable type 1 fusion machines are prevalent vaccine targets^20^.

Here we developed a modular self-assembling nanoparticle platform that allows for the plug-and-play display of trimeric viral glycoproteins on nanoparticle surfaces, utilizing the SpyTag:SpyCatcher system. We assessed this system with three prefusion (preF)-stabilized viral trimeric glycoproteins: respiratory syncytial virus fusion (RSV F) glycoprotein^21^, human parainfluenza virus type 3 fusion glycoprotein (PIV3 F)^22^, and SARS-CoV-2 spike glycoprotein^23,24^. SpyTag-coupled nanoparticles could be expressed with sufficient yield of soluble proteins from a mammalian expression system after the addition of nanoparticle surface glycans. The nanoparticle-formatted trimers exhibited improved antigenicity versus soluble trimers for apical epitopes, and we explicitly tested the immunogenicity for the nanoparticle-formatted trimeric antigen from SARS-CoV-2 in mice. Overall, protein antigens and nanoparticle scaffolds could be produced independently before conjugation, thereby expediting the otherwise generally cumbersome process of making and troubleshooting immunogens genetically fused to self-assembling nanoparticle subunits. Such a modular nanoparticle assembly platform may thus be a useful tool for plug-and-play screening of trimeric viral immunogens in a multivalent highly immunogenic context, and we provide proof-of-principle for increased immunogenicity of a nanoparticle-displayed SARS-CoV-2 spike.

## Results

### Expression of LuS- and ferritin-nanoparticle scaffolds with SpyTag requires the addition of an *N-*linked glycan

To construct a reliable plug-and-play platform for nanoparticle presentation of antigens, we chose *Aquifex aeolicus* lumazine synthase (LuS)^25^ and *Helicobacter pylori* ferritin^26^ as nanoparticle scaffolds with SpyTag:SpyCatcher conjugation system^15^ to display antigens on nanoparticle surface. The SpyTag:SpyCatcher system is highly specific and stable with an isopeptide bond and has been used for conjugation of antigens on nanoparticle surfaces^7,27^ (Fig. 1a). LuS and ferritin have served as scaffolds for nanoparticle immunogens in several clinical studies: for LuS see https://www.clinicaltrials.gov/ct2/show/NCT0369924128; for ferritin, see https://www.clinicaltrials.gov/ct2/show/NCT03547245^10,29,30^. The N terminus of both ferritin and LuS are exposed to the nanoparticle surface and are thus accessible for SpyTag or SpyCatcher attachment (Fig. 1b). The C terminus of LuS is also accessible on the nanoparticle surface and can be used to display purification tags. We designed mammalian expression constructs expressing fusion proteins of SpyTag or SpyCatcher with LuS or ferritin. The constructs included both His- and Strep-tags for purification purposes, along with a signal peptide for secretion of the expressed proteins into supernatant medium (Fig. 1b).

**Fig. 1.**
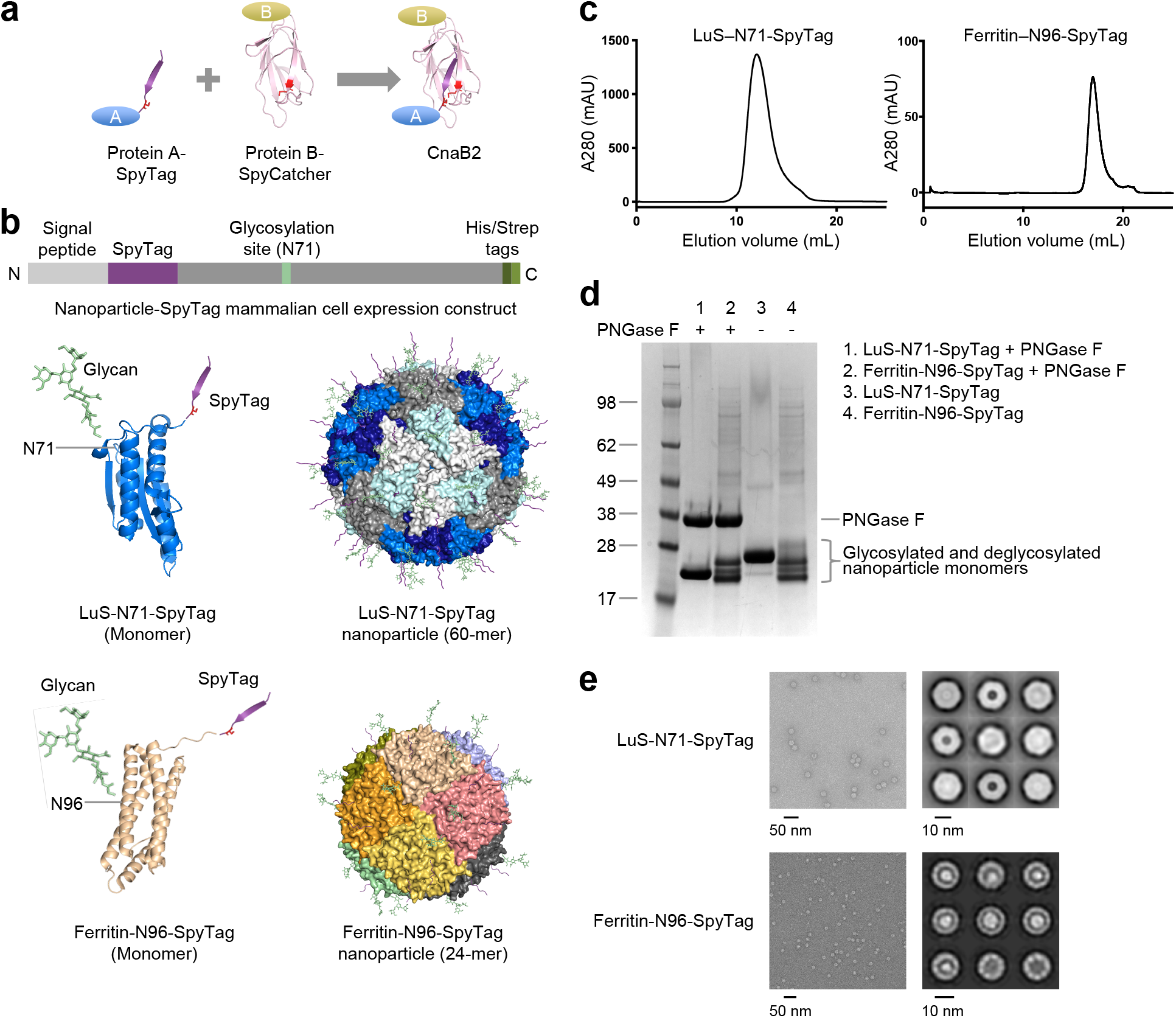
LuS- and ferritin-nanoparticle scaffolds with N-linked glycan and SpyTag express well as assembled nanoparticles in mammalian cells. (**a**) Schematic diagram showing the separate SpyTag and SpyCatcher to combine through an isopeptide bond as a means to covalently link molecules attached to SpyTag and molecules attached to SpyCatcher. (**b**) Design of expression constructs to produce activated nanoparticles with SpyTag in mammalian cells for conjugating antigens on the nanoparticle surface. Upper panel shows the DNA construct. A SpyTag was placed at the N-terminus of the nanoparticle sequence after the cleavable signal peptide. His and Strep tags were placed at the C-terminus of the LuS nanoparticle. An N-linked glycosylation site was engineered in the nanoparticle sequence to facilitate protein expression (see Table 1 and Supplementary Table S1 for more details). Lower panels show the expected structures of the LuS-N71-SpyTag and ferritin-N96-SpyTag monomers and assembled nanoparticles. Both glycan and SpyTag are expected to be on the nanoparticle surface. (**c**) Size exclusion chromatograms confirmed the correct sizes of the nanoparticles. The samples were loaded on a Superdex 200 Increase 10/300 GL column in PBS. Initial run of ferritin-96N-SpyTag nanoparticle revealed a tail of small molecular weight species; the chromatogram shown here is the re-run main peak. (**d**) SDS-PAGE of LuS-N71-SpyTag and ferritin-N96-SpyTag in the presence or absence of PNGase F. The position of PNGase F is marked. The multiple bands for ferritin are likely due to proteolytic cleavage and incomplete glycosylation (see text). (**e**) Negative stain EM images (left panels) and 2D class averages (right panels) of LuS-N71-SpyTag and ferritin-N96-SpyTag show the correct assembly of the purified nanoparticles with expected sizes.

Initial constructs yielded low levels of soluble proteins for the nanoparticle-SpyTag or SpyCatcher fusion proteins. To improve protein solubility and expression, we added glycans to the surface of the nanoparticles, designing a panel of LuS and ferritin constructs with SpyTag and SpyCatcher (Table 1 and Supplementary Table S1). For LuS constructs, we added a glycosylation site at position 71 (PDB 1HQK numbering). For ferritin constructs, two potential glycosylation sites (96 and 148) were tested. The addition of *N*-linked glycosylation sites facilitated expression of soluble nanoparticles in the cell culture supernatant. Three of the constructs produced appreciable yields of well-assembled nanoparticles, LuS with N71 and SpyTag at N-terminus (hereafter referred to as LuS-N71-SpyTag), ferritin with N96 and SpyTag, and ferritin S148 (glycan at N146) and SpyTag (Table 1). Of the two ferritin constructs, the ferritin with N96 and SpyTag had a higher yield and was chosen for further studies (hereafter referred to as ferritin-N96-SpyTag). Size exclusion chromatography (SEC) and electron microscopy (EM) analyses indicated that LuS-N71-SpyTag formed a homogeneous nanoparticle population in solution (Fig. 1c,d). The ferritin-N96-SpyTag sample comprised mainly intact nanoparticles with some minor unassembled species (Fig. 1c,d). Negative-stain electron microscopy (EM) images indicated both nanoparticles to be well-assembled with expected sizes^25,26^ (Fig. 1d). Two-dimensional class average revealed more detailed structural features of the nanoparticles, which were consistent with previously published structures of the two nanoparticles. These data indicated the ferritin and LuS nanoparticles were compatible with the SpyTag and glycosylation site addition. These alterations were well tolerated, allowing for robust nanoparticle assembly. To verify the glycosylation of LuS- and ferritin-SpyTag nanoparticles, we performed PNGase F digestion and checked for glycan cleavage through SDS-PAGE (Fig. 1e). Both nanoparticles showed a band shift in the presence of PNGase F, indicating the presence of *N-*liked glycan on the nanoparticles and its removal by the amidase digestion. While the glycan cleavage in LuS-N71-SpyTag is distinct, it is less apparent in ferritin-N96-SpyTag, likely due to incomplete glycosylation of ferritin-N96-SpyTag and multiple bands of ferritin on SDS-PAGE. Ferritin has been observed to exhibit a single band on SDS-PAGE in some studies^2^ but multiple bands in others^16,31^, presumably due to protease cleavage at the C terminus or incomplete glycosylation. However, these different sized ferritin molecules assembled correctly as nanoparticles with expected dimensions as indicated by SEC and EM (Fig. 1c,e).

**Table 1.**
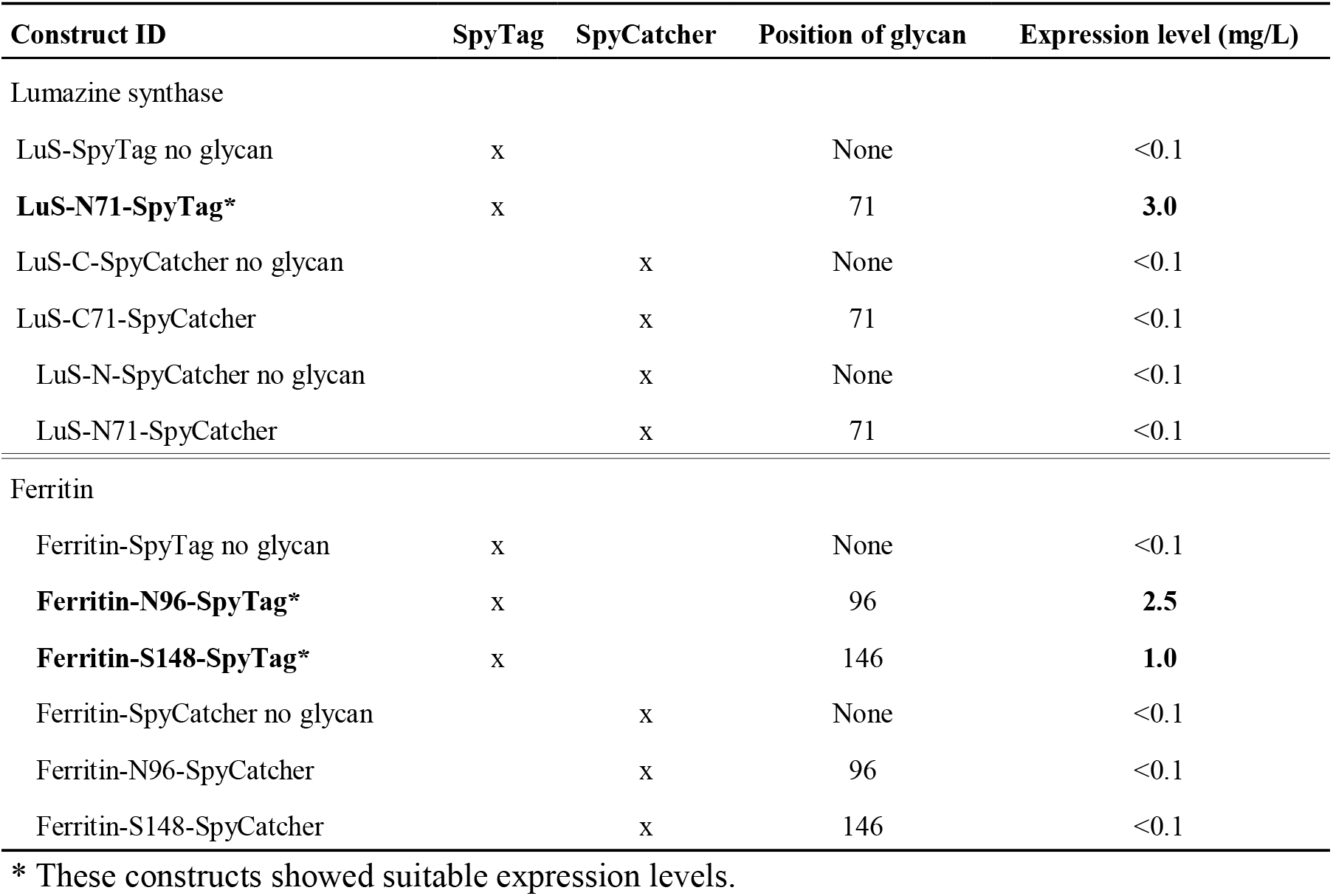
LuS- and ferritin-nanoparticles with SpyTag require the addition of *N*-linked glycans for expression.

### Conjugation of RSV F-SpyCatcher to LuS-N71-SpyTag displays prefusion RSV F ‘DS2’-trimers homogeneously on the surface of the LuS-N71-Spylinked-RSV F nanoparticle

With no effective licensed vaccine against respiratory syncytial virus (RSV), the causative agent for a prevalent childhood disease that results in approximately 60,000 hospitalizations and 10,000 annual deaths in the US, developing an immunogen capable of eliciting protection is of paramount importance^32^. Previous efforts in developing a vaccine capable of eliciting protective antibodies have resulted in the identification of RSV trimers stabilized in its prefusion conformation, RSV F DS-Cav1 (DS-Cav1), and RSV F DS2 (DS2)^21,33^. DS2 was shown to elicit higher RSV neutralization responses than DS-Cav1^33^. With this as our motivation and as a test case for our nanoparticle SpyTag:SpyCatcher system, we investigated the feasibility of displaying DS2 in the context of nanoparticle immunogens.

We prepared DS2 coupled to SpyCatcher (hereafter referred to as RSV F-SpyCatcher) by genetic engineering to append SpyCatcher to the C-terminus of RSV F after a 3 residue (GSG) linker (Supplementary Table S1). After expression and purification, we conjugated the purified RSV F-SpyCatcher to the purified 60-mer LuS-N71-SpyTag nanoparticle (Fig. 2a). SEC profiles of the two components revealed that LuS-N71-SpyTag eluted around 13 mL and RSV F-SpyCatcher eluted near 15 mL on Superdex 200 Increase 10/300 column (GE Health Sciences) (Fig. 2b). The conjugated LuS-N71-SpyLinked-RSV F-SpyCatcher nanoparticle (LuS-N71-SpyLinked-RSV F) eluted in a new peak at ~10 mL by SEC (Fig. 2b). SDS-PAGE showed the appearance of species of larger molecular weight of ~90 kDa in the conjugation mixture, followed by bands of residual LuS-N71-SpyTag monomer and RSV F-SpyCatcher components at 20 kDa and 60 kDa, respectively (Fig. 2c), confirming the success of the conjugation reaction. To estimate the conjugation efficiency, we measured the intensity of each band on the SDS-PAGE gel image of the conjugated nanoparticle product (Fig. 2c), as a surrogate of mass for each component.

**Fig. 2.**
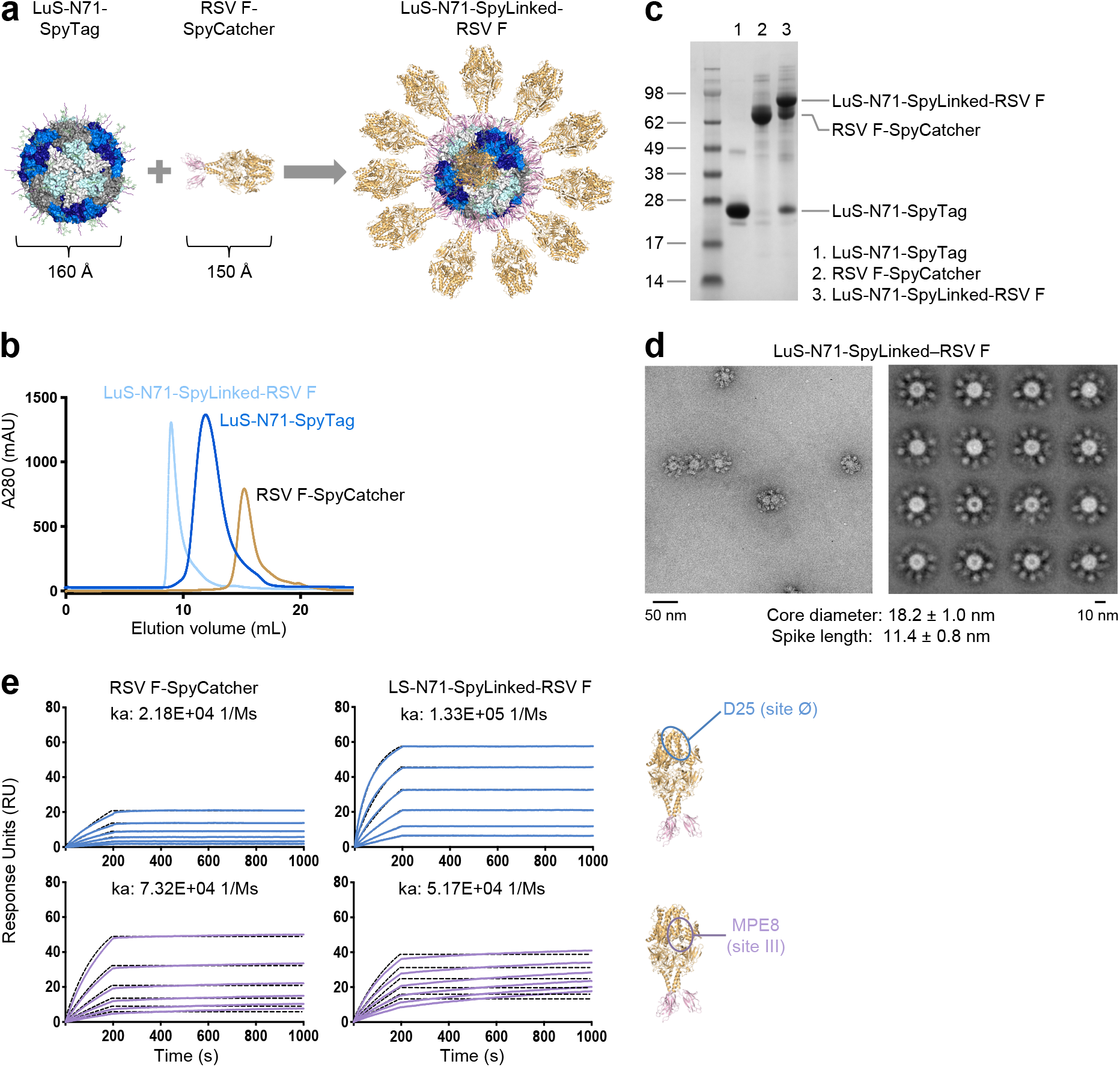
Conjugation of RSV F-SpyCatcher to LuS-SpyTag displays prefusion RSV F trimer homogenously on the surface of the LuS-N71-SpyLinked-RSV F nanoparticle. (**a**) Schematic diagram showing conjugation of SpyTag-coupled LuS to SpyCatcher-coupled RSV prefusion F trimer to make LuS-N71-SpyLinked-RSV F nanoparticle. (**b**) SEC profiles of LuS-N71-SpyTag, RSV F-SpyCatcher, and the conjugated product LuS-N71-SpyLinked-RSV F on a Superdex 200 Increase 10/300 GL column in PBS. (**c**) SDS-PAGE of LuS-N71-SpyTag (lane 1), RSV F-SpyCatcher (lane 2), and the conjugated LuS-N71-SpyLinked-RSV F nanoparticle product (lane 3), in the presence of DTT. (**d**) Negative stain EM images of the LuS-N71-SpyLinked-RSV F nanoparticle after SEC purification, showing (left panel) a representative micrograph and (right panel) the 2D class averages. (**e**) Surface plasmon resonance of RSV F-SpyCatcher and LuS-N71-SpyLinked-RSV F nanoparticle with prefusion-specific D25 IgG (site Ø) and MPE8 IgG (site III), with IgG coupled to chip and nanoparticle in solution. A concentration series from 200 nM to 1.56 nM of RSV F either as trimer (left) or couped to nanoparticle (right) was measured; k_a_ values are provided as these have been found to correlate with immunogenicity^3^.

Taking into consideration the molecular weight of each component, we calculated the molar ratio of each component to total protein in the sample. We estimated 67% of all the LuS nanoparticle subunit was conjugated to RSV F trimer. To verify particle integrity after conjugation, we performed negative stain EM following SEC purification. LuS-N71-SpyTag conjugated with RSV F-SpyCatcher efficiently produced uniform particles with a core diameter of 18.2 ± 1.0 nm decorated with trimer spike of 11.4 ± 0.8 nm in length (Fig. 2d). We then confirmed the prefusion state of the LuS-N71-SpyLinked RSV F nanoparticle through surface plasmon resonance using RSV prefusion F specific antibodies D25 (site Ø) and MPE8 (site III) (Fig. 2e)^21^. Notably, RSV F on nanoparticles showed an enhanced on-rate to the apex-targeting D25 antibody and reduced on-rate to the equatorial targeting MPE8 versus trimeric RSV F, a crucial antigenic characteristic signifying appropriate nanoparticle display^3^.

### Conjugation of RSV prefusion F-SpyCatcher to ferritin-SpyTag produces uniform ferritin-RSV F nanoparticles

Having produced successfully the LuS-N71-SpyLinked-RSV F nanoparticle, we next set out to conjugate the 24-mer ferritin-N96-SpyTag with RSV F-SpyCatcher in the same manner (Fig. 3a). SEC of ferritin-N96-SpyTag nanoparticle showed a peak at around 16-17 mL, slightly slower than RSV F-SpyCatcher (Fig. 3b). Negative stain EM revealed that ferritin-N96-SpyTag formed nanoparticle of the expected size (Fig. 1c). The conjugation mixture of ferritin-N96-SpyTag with RSV F-SpyCatcher exhibited a peak at ~10 mL (void volume of the SEC column), suggesting successful formation of the conjugation product (referred to as ferritin-RSV F) (Fig. 3b). SDS-PAGE demonstrated the appearance of a new band at ~90 kDa, the expected size of ferritin-N96-SpyLinked-RSV F nanoparticle, with residual ferritin-N96-SpyTag at around 20 kDa (Fig. 3c). Using the same method as for LuS-N71-SpyLinked-RSV F above, we estimated 85% of all the ferritin nanoparticle subunit was conjugated to RSV F trimer. To confirm the formation of ferritin-N96-SpyLinked-RSV F nanoparticle, we performed negative stain EM, which showed well-formed nanoparticles with the expected size and shape, displaying trimer spikes around the ferritin nanoparticle (Fig. 3d).

**Fig. 3.**
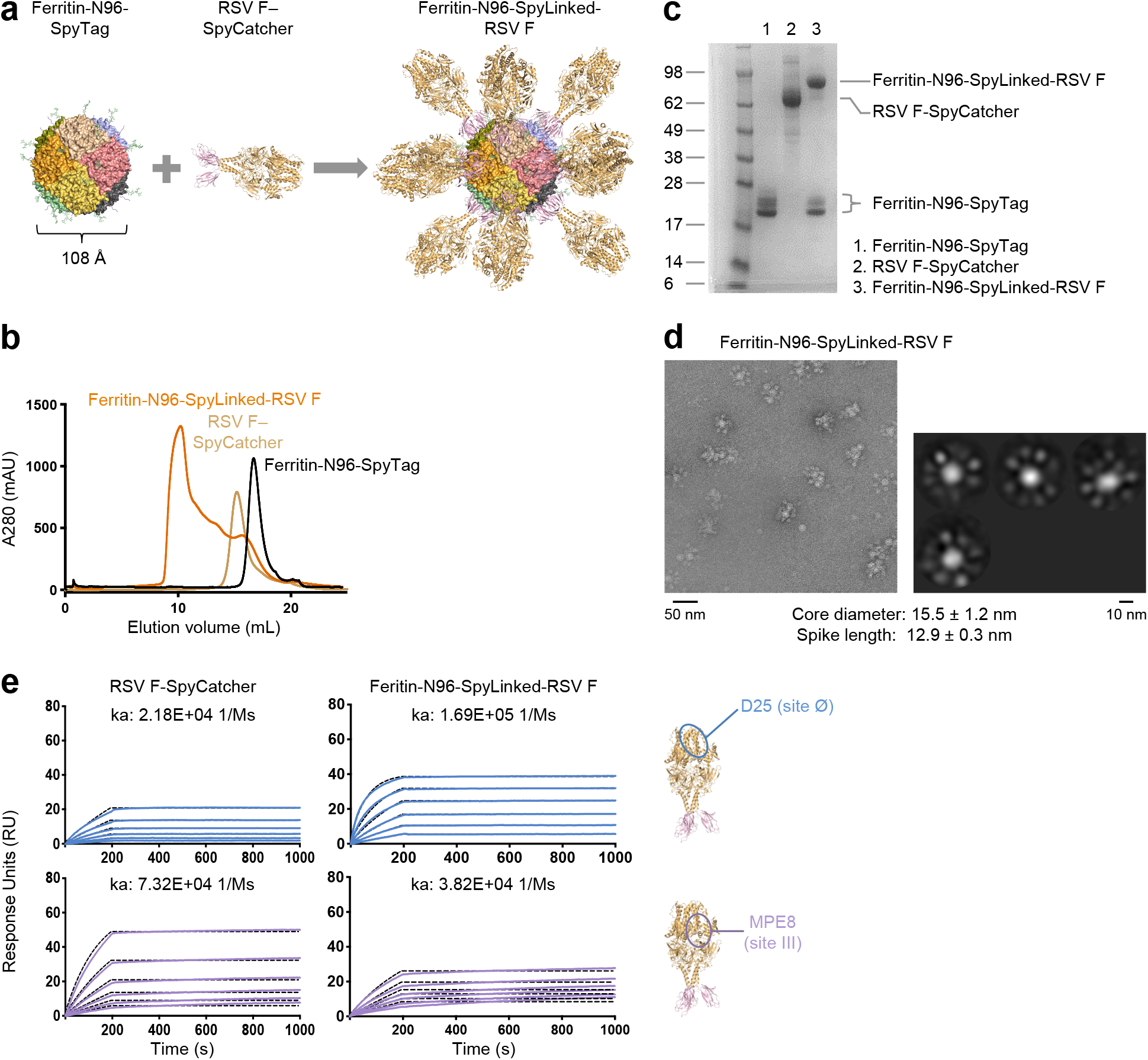
Conjugation of RSV prefusion F-SpyCatcher to ferritin-SpyTag produces uniform ferritin-RSV F nanoparticles. (**a**) Schematic diagram showing the conjugation process of ferritin-N96-SpyTag and RSV F– SpyCatcher to make ferritin-N96-SpyLinked-RSV F nanoparticle. (**b**) SEC profiles of ferritin-N96-SpyTag, RSV F-SpyCatcher, and the conjugation reaction mixture on a Superdex 200 Increase 10/300 GL column in PBS. (**c**) SDS-PAGE of ferritin-N96-SpyTag (lane 1), RSV F-SpyCatcher (lane 2), and the conjugated ferritin-N96-SpyLinked-RSV prefusion F SpyCatcher nanoparticle product (lane 3), in the presence of DTT. Ferritin exhibited multiple bands due to proteolytic cleavage and incomplete glycosylation (see text). (**d**) Negative stain EM images of the ferritin-RSV F nanoparticle after SEC purification, showing (left panel) a representative micrograph and (right panel) the 2D class averages. (**e**) SPR of RSV F-SpyCatcher and Ferritin-N96-SpyLinked-RSV F nanoparticle with prefusion-specific D25 IgG (site Ø) and MPE8 IgG (site III) using immobilized IgG on sensor chip with nanoparticle and trimer in solution. A concentration series from 200 nM to 1.56 nM of RSV F either as trimer (left) or coupled to nanoparticle (right) was measured; k_a_ values are provided.

To verify the conserved prefusion state of the conjugated RSV F trimer, we measured the binding of ferritin-N96-SpyLinked-RSV F to D25 and MPE8 IgGs through SPR (Fig. 3e). Importantly, we observed the on-rate to increase for D25, which recognizes an epitope at the trimer apex, but the on-rate to decrease for MPE8, which recognizes an equatorial epitope on the trimer, similar to the observation for LuS-N71-SpyLinked-RSV F.

### Displaying PIV3 F glycoprotein trimer on LuS nanoparticle via SpyTag:SpyCatcher conjugation improves antibody binding to the trimer apex

To demonstrate the plug-and-play versatility of the SpyTag:SpyCatcher nanoparticle system, we produced PIV3 F glycoprotein trimer^34^ as a fusion protein with SpyCatcher at the C terminus and conjugated with the LuS-N71-SpyTag nanoparticle (Fig. 4, a-c, Supplementary Table S1), similar to that described above for the conjugation of RSV F-SpyCatcher. PIV3 is a prevalent human parainfluenza virus that causes respiratory illnesses, especially in infants and young children^35,36^. The conjugation mixture of PIV3 F-SpyCatcher and LuS-N71-SpyTag was loaded onto the SEC column to purify the conjugated nanoparticle product LuS-N71-SpyLinked-PIV3 F from unconjugated nanoparticles and the PIV3 F SpyCatcher trimer (Fig. 4b). SDS-PAGE analysis revealed that the conjugated product had the expected molecular weight and there was not unconjugated PIV3 F-SpyCatcher in the conjugation mixture (Fig. 4c). Using the same method as for LuS-N71-SpyLinked-RSV F in the previous section, we estimated 68% of all the LuS nanoparticle subunit was conjugated to PIV3 F trimer. The PIV3 F conjugated nanoparticle was further verified through negative stain EM, which showed well defined trimer spikes decorating the LuS nanoparticle with the expected size (Fig. 4d).

**Fig. 4.**
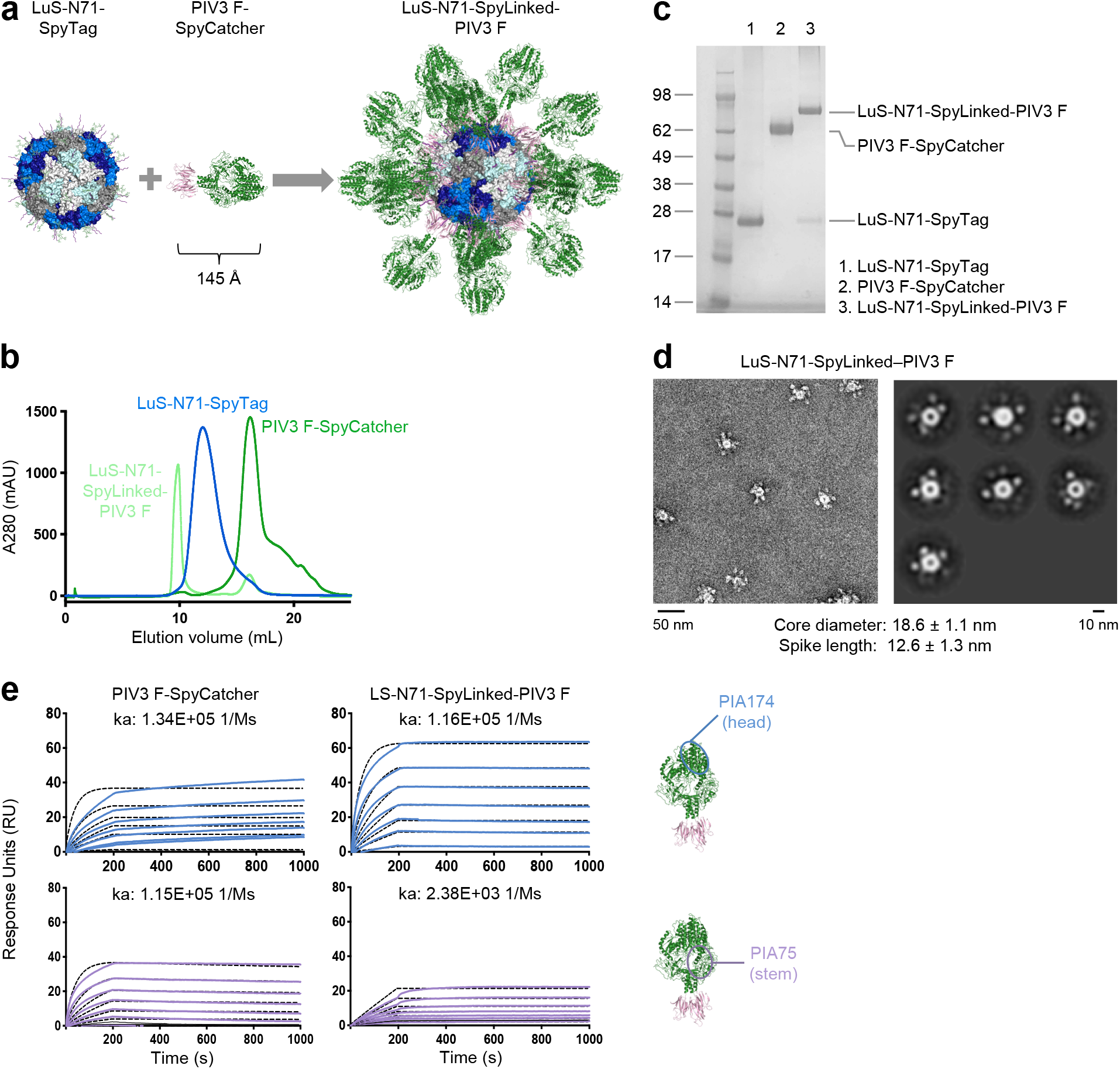
Conjugation of PIV3 F-SpyCatcher to LuS-SpyTag displays prefusion PIV3 F trimer homogenously on the surface of the LuS-N71-SpyLinked-PIV3 F nanoparticle. (**a**) Schematic of the conjugation between LuS-N71-SpyTag and PIV3 F-SpyCatchert to produce LuS-N71-SpyLinked-PIV3 F nanoparticle (**b**) SEC profiles of PIV3 F-SpyCatcher, LuS-SpyTag, and the conjugated product LuS-N71-SpyLinked-PIV3 F on a Superdex 200 Increase 10/300 GL in PBS. (**c**) SDS-PAGE of LuS-N71 (lane 1), PIV3 F-SpyCatcher (lane 2), and LuS-N71-SpyLinked-PIV3 F conjugation mixture (lanes 3) in the presence of DTT. (**d**) Negative stain EM of LuS-N71-SpyLinked-PIV3 following SEC showing a representative micrograph (left panel) and 2D class averages (right panel). (**e**) SPR measurements of PIV3 F-SpyCatcher and LuS-N71-SpyLinked-PIV3 F were performed using IgG coupled chips with nanoparticle and timer in solution. A concentration series from 200 nM to 1.56 nM of PIV3 F either as trimer (left) or coupled to nanoparticle (right) was measured; k_a_ values are provided.

Having produced nanoparticles of LuS-N71-SpyTag conjugated with PIV3 F-SpyCatcher, we next evaluated binding of LuS-N71-SpyLinked-PIV3 F with antibodies PIA174 and PIA75^34^, using SPR (Fig. 4e). The head-targeting antibody PIA174 showed an improved binding to LuS-N71-SpyLinked-PIV3 F relative to its binding to PIV3 F-SpyCatcher. The stem-targeting antibody PIA75, however, showed decreased binding to LuS-N71-SpyLinked-PIV3 F compared with PIV3 F-SpyCatcher.

### Conjugation of SARS-CoV-2 spike trimer to LuS nanoparticle via SpyTag:SpyCatcher displays the spike trimers homogeneously on the nanoparticle surface

Severe acute respiratory syndrome coronavirus 2 (SARS-CoV-2) caused the COVID-19 pandemic that is ongoing worldwide^37^. An effective vaccine against SARS-CoV-2 and related coronaviruses is urgently needed. The SARS-CoV-2 spike glycoprotein trimer mediates virus-cell membrane fusion and thus a target for vaccine development^23,24^. To test the versatility of our plug-and-play SpyTag:SpyCatcher nanoparticle system, we expressed and purified SARS-CoV-2 spike fused with a C-terminal SpyCatcher and conjugated to the LuS-N71-SpyTag nanoparticle (Fig. 5, a-c, Supplementary Table S1). For this construct, we used the prefusion stabilized version of spike developed by McLellan and colleagues^24^, which included GSAS and PP mutations and the T4 phage fibritin trimerization domain along with a single-chain Fc tag as described by Zhou and colleagues^38^.

**Fig. 5.**
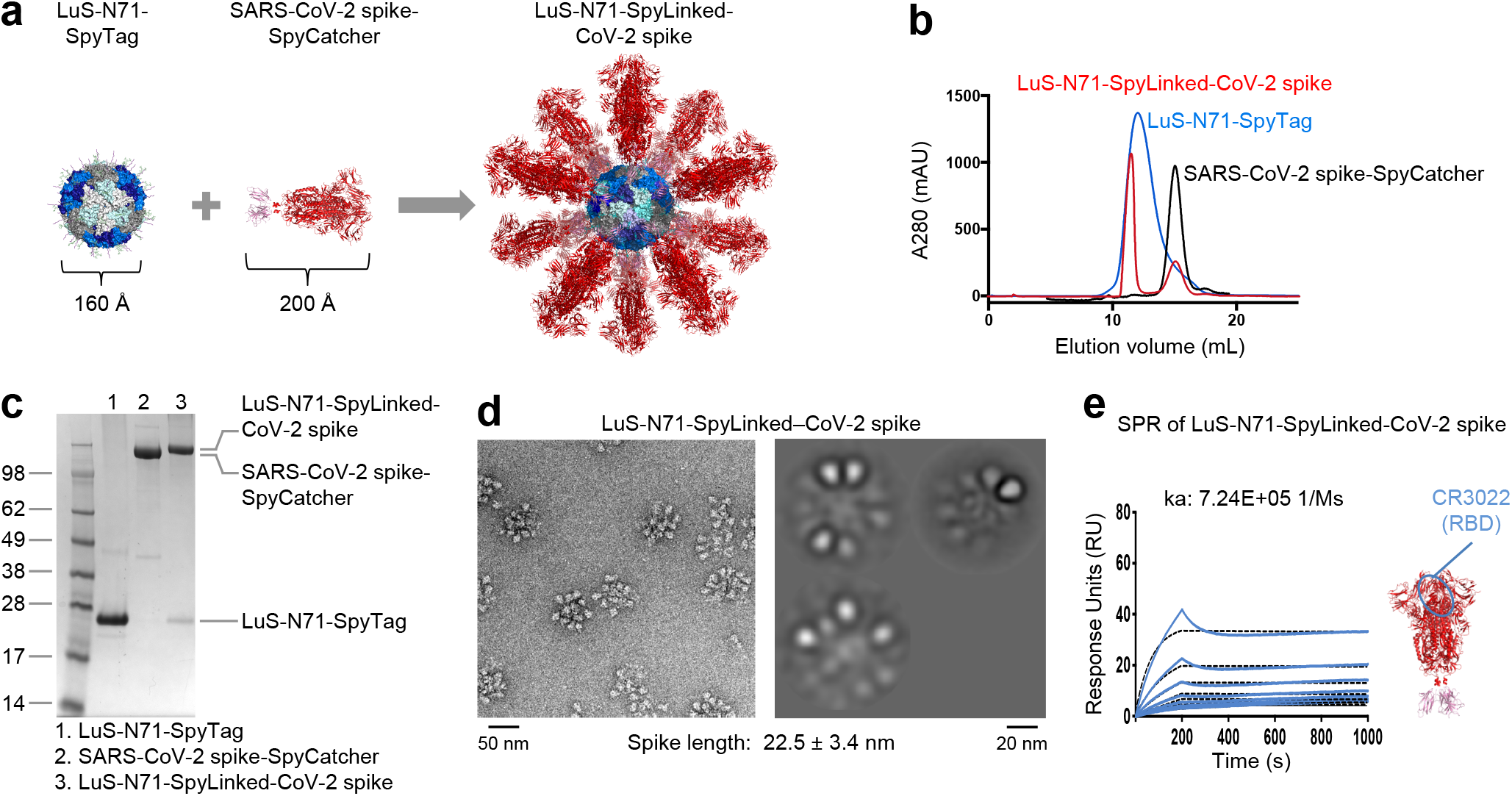
Conjugation of SARS-CoV-2 spike trimer to LuS-SpyTag displays SARS-CoV-2 spike trimer on the surface of the LuS-N71-SpyLinked-CoV-2 spike nanoparticle. (**a**) Schematic diagram showing conjugation of SpyTag-coupled LuS to SpyCatcher-coupled SARS-CoV-2 spike trimer to make LuS-N71-SpyLinked-CoV-2 spike nanoparticle. (**b**) SEC profiles of LuS-N71-SpyTag, SARS-CoV-2 spike-SpyCatcher, and the conjugated product LuS-N71-SpyLinked-CoV-2 spike on a Superdex 200 Increase 10/300 GL column in PBS. (**c**) SDS-PAGE of LuS-N71-SpyTag (lane 1), SARS-CoV-2 spike-SpyCatcher (lane 2), and the conjugation mixture of LuS-N71-SpyTag with SARS-CoV-2 spike-SpyCatcher (lane 3) in the presence of DTT. The conjugation mixture (lane 3) shows the conjugated LuS-N71-SpyLinked-CoV-2 spike nanoparticle with minor excess of LuS-N71-SpyTag. (**d**) Negative stain EM of the LuS-N71-SpyLinked-CoV-2 spike nanoparticle after SEC purification showing representative micrographs (left panel) and 2D class average (right panel). (**e**) SPR response curves for LuS-N71-SpyLinked-CoV-2 spike nanoparticle binding with RBD-targeting antibody CR3022 IgG, with IgG coupled to chip and nanoparticle in solution. Because SARS-CoV-2 spike-SpyCatcher showed non-specific binding only the coupled nanoparticle is shown. A series of nanoparticle concentrations was analyzed in which the concentration of spike coupled to the nanoparticle ranged from 200 nM to 1.56 nM. Observed k_a_ value provided.

The conjugation mixture was loaded onto an SEC column to purify the conjugated nanoparticle product LuS-N71-SpyLinked-CoV spike from unconjugated LuS-N71-SpyTag and SARS-CoV-2 spike-SpyCatcher (Fig. 5b). SDS-PAGE analysis revealed the conjugated product to have the expected molecular weight, and unconjugated spike-SpyCatcher was not observed after conjugation (Fig. 5c). Using the same method as for LuS-N71-SpyLinked-RSV F, we estimated 91% of all the LuS nanoparticle subunit was conjugated to the spike trimer. Negative stain EM showed LuS-N71-SpyLinked-CoV-2 spike nanoparticle to exhibit the expected size with spike trimers displaying on the LuS nanoparticle surface (Fig. 5d). SPR measurements showed LuS-N71-SpyLinked-SARS-CoV-2 Spike to bind to CR3022^39,40^, an antibody targeting the receptor-binding domain (RBD), indicating successful nanoparticle presentation of the spike trimer using the LuS-SpyTag:SpyCatcher system.

### LuS-N71-SpyLinked-nanoparticle display increases potential of SARS-CoV-2 spike to elicit neutralizing antibodies

To assess immunogenicity, we injected mice with the LuS-N71-SpyLinked-CoV-2 spike nanoparticle or spike trimers (stabilized by 2P mutation)^24,41^, or mock (LuS-N71-SpyTag) nanoparticles at weeks 0 and 3 (Fig. 6a). Serum samples were collected two weeks after each immunization. After the first immunization, at the lowest immunogen dose of 0.08 μg, spike nanoparticle-immune sera exhibited an anti-SARS-CoV-2 spike ELISA geometric mean titer of 5,116, whereas only 1 out of 10 trimeric spike-immunized sera exhibited a measurable titer (Fig. 6b); after a second immunization, titers for the spike nanoparticle-immune sera increased substantially, by approximately 25-fold. Immunizations with higher doses of spike nanoparticle (0.4 and 2.0 μg) increased titers more incrementally, both at week 2 and at week 5. By contrast, increases in dose of the spike trimer raised ELISA titers more substantially, with two of the mice in the 2.0 μg spike-trimer immune sera reaching the assay upper limit of detection with a titer of 1,638,400 (Fig. 6b).

**Fig. 6.**
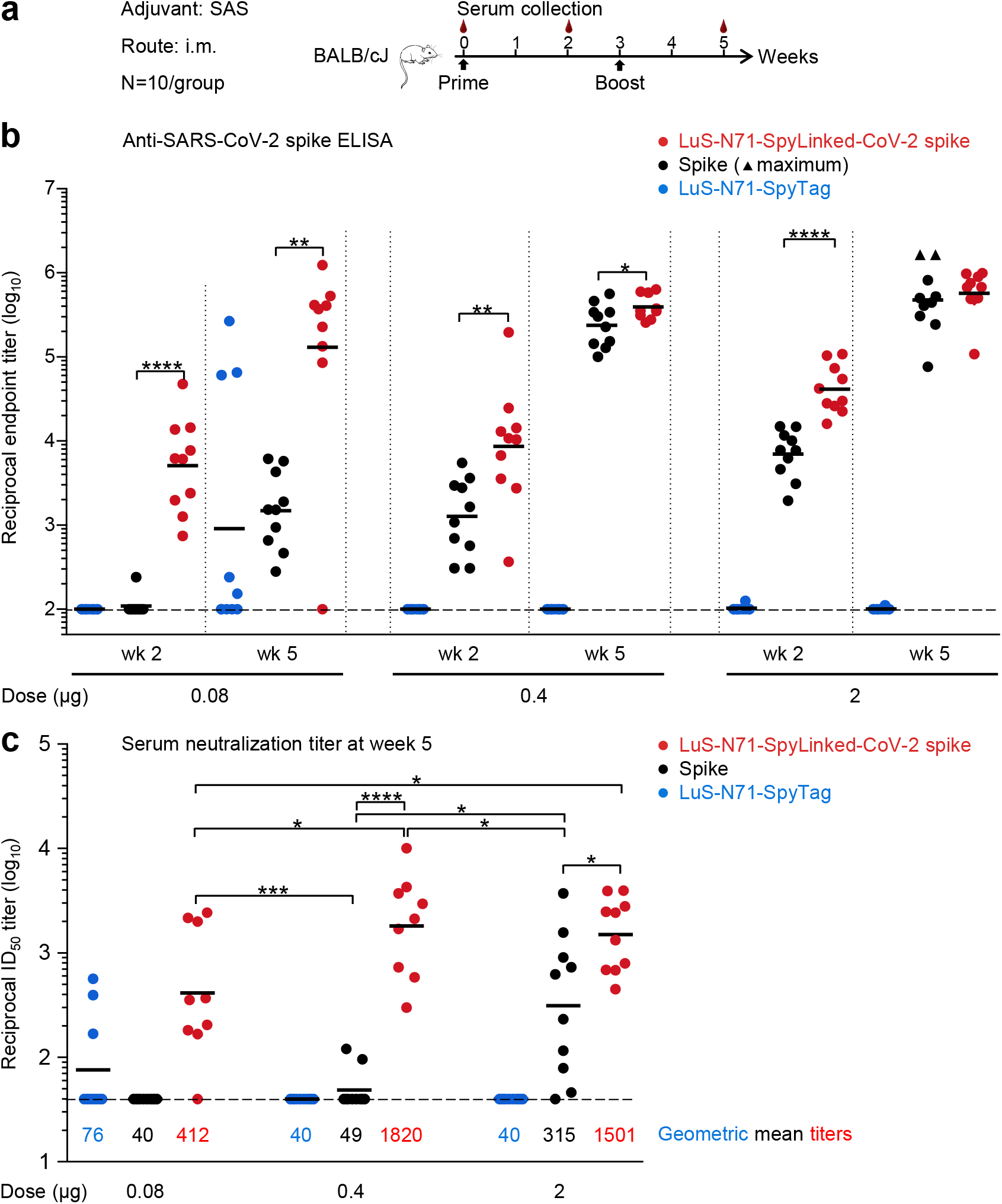
Immunogenicity of LuS-N71-SpyLinked-CoV-2 spike. (**a**) Schematic immunization procedures for SARS-CoV-2 spike immunogens. (**b**) Serum assessment of anti-SARS-CoV-2 spike ELISA titers. Immunization groups are color-coded. Vertical dotted lines separate immunogen dose groups and weeks post prime. Starting reciprocal serum dilution (100) is indicated with a horizontal dashed line. ELISA titer from each animal is shown as an individual dot. Triangle-shape dot provided for ELISA titers at assay maximum. Geometric means indicated by black horizontal lines. Note that the three animals immunized with 0.08 μg LuS-N71-SpyTag, which showed high ELISA titers at week 5, were the same three animals of this control group that showed detectable neutralization. (**c**) Neutralization titer from each animal at week 5 is shown as an individual dot, and geometric means are indicated by black horizontal lines with values provided for each group. Immunization groups are color-coded as in panel **b**. Limit of detection (titer = 40) indicated with a horizontal dashed line. *P* values determined by two-tailed Mann-Whitney tests. * indicates *P* ≤ 0.05, ** indicates *P* ≤ 0.01, *** indicates *P* ≤ 0.001 and **** indicates *P* ≤ 0.0001.

Importantly, pseudovirus neutralization assays revealed the LuS-N71-SpyLinked-CoV-2 spike nanoparticle to elicit potent neutralization responses with geometric mean ID_50_ titers of 413, 1820, and 1501 for immunization doses of 0.08, 0.4, and 2 μg, respectively (Fig. 6c). In comparison, two doses of trimeric spike elicited neutralization titers at the 0.4 and 2 μg doses with a geometric mean ID_50_ of 49 and 315, respectively, with no measurable neutralization at the 0.08 μg dose. In essence, 0.08 μg of spike nanoparticle elicited a neutralization response that was higher, though statistically indistinguishable from 2 μg of trimeric spike. This indicated ~25-fold higher immunogenicity on a weight-by-weight basis for the spike nanoparticle versus spike alone, suggesting a substantial “dose-sparing” effect. Overall, presentation of the SARS-CoV-2 spike on the LuS nanoparticle surface significantly improved its immunogenicity and required a lower immunogen dose to elicit potent neutralization responses compared with the trimeric form.

## Discussion

Nanoparticle-based immunogens can induce potent neutralizing antibodies^2,3,42^ and thus may be promising vaccine candidates. To develop nanoparticle vaccine immunogens, rapid and efficient methods would help produce nanoparticle scaffolds that can be mixed and matched with different immunogens. Previous efforts utilizing the spontaneous isopeptide bond formation with the SpyTag:SpyCatcher system for nanoparticle surface display of immunogens^11–16^ have proven the versatility of this system for antigen display. However, none of these previously published reports utilized mammalian expression allowing for post-translational modifications, such as *N*-linked glycosylation. Here, we describe two nanoparticle platforms, lumazine synthase and ferritin, for the display of trimeric viral protein immunogens using the SpyTag:SpyCatcher system. By adding *N-*linked glycosylation sites to nanoparticle monomers, we were able to produce SpyTag-coupled nanoparticles using mammalian cell culture.

LuS and ferritin nanoparticle platforms vary in the number of molecules displayed on the surface. LuS-N71-SpyTag contains 60 SpyTags whereas ferritin-N96-SpyTag has 24 displayed on surface, available for SpyCatcher-carrying molecules to couple to. Both platforms showed efficient conjugation of trimeric immunogens and formed nanoparticle rapidly under physiological conditions for RSV F, PIV3 F and SARS-CoV-2 spike trimers. One advantage of the LuS nanoparticle is the high efficiency of its particle assembly. The glycosylated LuS-N71-SpyTag assembled into a homogenous particle that exhibited a single peak in size exclusion chromatography.

To demonstrate the versatility of our SpyTag-displaying nanoparticles in immunogen development, we conjugated them to three viral antigens of vaccine interest, the DS2-preF stabilized RSV F^33^, a DS2-stabilized version of PIV3 F^34^, and the 2P-stabilized version of SARS-CoV-2 spike^24^. In each of these, we appended the SpyCatcher after the ‘foldon’ heterologous trimeric stabilization motif^43^. Conjugation of SpyTag-nanoparticles with SpyCatcher-coupled RSV F, PIV3 F and SARS-CoV-2 spike trimers resulted in proper particle assembly. In all three cases, we observed high conjugation efficiency.

Our antigenicity analyses indicate that presentation of trimeric antigens from viral pathogens on self-assembling nanoparticles needs to take into consideration the accessibility of the antigenic epitopes. When a trimer protein is conjugated to a nanoparticle, such as LuS or ferritin in this study, the trimer molecules are densely displayed on the nanoparticle surface. As a result of the dense display, the epitopes near the nanoparticle surface, such as those at the stem region of the trimers in this study, are less accessible to antibodies than the epitopes on the apex of trimer molecules. Consequently, we observed an increased level of antibody binding to epitopes on the apex and a decreased level of antibody binding to epitopes on the equatorial or stem region of RSV F and PIV3 F trimer molecules (Figs. 2e, 3e and 4e).

The increased antigenicity of the apical epitopes on the trimer conjugated to nanoparticles is expected to yield increased immunogenicity – especially at lower dose, and we provide proof-of-principle for this with murine immunization studies with LuS-N71-SpyLinked-CoV-2 spike as compared to soluble spike. As observed in prior studies^2,3^, nanoparticle immunogens elicited stronger immune responses than the corresponding trimers at low immunogen doses: at the 0.08 μg dose after two immunizations, spike nanoparticle elicited neutralization response with ID_50_ of 413, whereas trimeric spike elicited an equivalent neutralization titer only at the 25-fold higher dose of 2 μg. At 0.4 μg, spike nanoparticle elicited ~37-fold higher ID_50_ than trimeric spike.

However, at a high dose of 2 μg, spike nanoparticle-elicited neutralization response appeared to plateau – at a level ~5-fold higher in neutralization titer than the trimeric immunogen. Similar increases in immunogenicity and with dose-sparing have been recently reported for nanoparticles incorporating the receptor-binding domain (RBD) of the spike^44^. Overall, multivalent presentation of trimeric antigens on nanoparticle can significantly improve their immunogenicity, allowing for elicitation of potent immune responses at a relatively low immunogen dose. Our SpyTag:SpyCatcher system provides a versatile platform for preparation of such nanoparticle immunogens from trimeric antigens.

It will be interesting to see if the plug-and-display technology described here will allow for the incorporation of different molecules on multiple nanoparticles. Such molecules could include not only trimeric viral immunogens, but immunostimulatory components, or molecules targeting antigen presenting cells. Thus, the LuS- and ferritin-SpyTag displaying nanoparticles described here may be amendable to mix-and-match display of immunogens and of immunostimulatory or targeting components.

## Materials and Methods

### Protein production and purification

The amino acid sequences of protein expression constructs are listed in Supplementary Table S1. For protein expression, 3 ml of Turbo293 transfection reagent (Speed BioSystems) was mixed with 50 ml Opti-MEM medium (Life Technology) and incubated at room temperature (RT) for 5 min. 1 mg plasmid DNAs was mixed with 50 ml of Opti-MEM medium in a separate tube, and the mixture added to the Turbo293 Opti-MEM mixture. The transfection mixture was incubated for 15 min at RT then added to 800 ml of Expi293 cells (Life Technology) at 2.5 million cells/ml. The transfected cells were incubated overnight in a shaker incubator at 9% CO_2_, 37 °C, and 120 rpm. On the second day, about 100 ml of Expi293 expression medium was added. On day 5 post transfection, supernatants were harvested, filtered. Proteins were purified from the supernatant using Ni-NTA and strep chromatography. SARS-CoV-2 spike-SpyCatcher was expressed as a fusion protein with a single-chain Fc purification tag and purified using Protein A chromatography. SARS-CoV-2 spike-SpyCatcher protein was cleaved off from Protein A column by HRV3C protease. All proteins were further purified by size exclusion chromatography on Superdex 200 Increase 10/300 GL in PBS.

### LuS- and ferritin-SpyTag conjugations

A 1:1 molar ratio of LuS- or ferritin-SpyTag and immunogen-SpyCatcher components were combined and incubated at ambient temperature for approximately 3 hours, followed by size exclusion column on Superdex200 Increase 10/300 GL in PBS to separate conjugated products from residual components. The conjugated nanoparticle product was then run through SDS-PAGE to verify conjugation and analyzed by negative-stain EM.

### Negative-stain electron microscopy (EM)

Samples were diluted to 0.02-0.05 mg/ml with a buffer containing 10 mM HEPES, pH 7, and 150 mM NaCl. A 4.7-μl drop of the diluted sample was applied to a glow-discharged carbon-coated copper grid for approximately 15 s. The drop was removed using blotting paper, and the grid was washed three times with 4.7-μl drops of the same buffer. Adsorbed proteins were negatively stained by applying consecutively three 4.7-μl drops of 0.75% uranyl formate and removing each drop with filer paper. Micrographs were collected using SerialEM^45^ on an FEI Tecnai T20 electron microscope operated at 200 kV and equipped with an Eagle CCD camera or using EPU on a ThermoFisher Talos F200C electron microscope operated at 200 kV and equipped with a Ceta CCD camera. The pixel size was 0.44 and 0.25 nm for Tecnai T20 and Talos F200C, respectively. Particles were picked automatically using in-house written software (Y.T., unpublished). Reference-free 2D classification was performed with Relion 1.4^46^ and SPIDER^47^. The dimensions of VLP cores and spikes were measured with e2display.py program from EMAN2.1 software package^48^ using a representative micrograph (LuS-N71-SpyLinked–CoV-2 S) or 2D class average images (all other VLPs). For CoV-2 nanoparticles we observed increased structural content at low pH, whereas the other micrographs were collected at physiological pH.

### Surface plasmon resonance (SPR)

To prepare the chips (GE Healthcare Life Sciences CM5 chips), antibody IgG were immobilized onto the chip by amine coupling to ~100-1000 response units (RU) depending the level of binding of each trimer and nanoparticle pair to antibodies. To measure binding of SpyCatcher proteins and and nanoparticles, a dilution series of SpyCatcher-linked proteins and nanoparticles were flowed through the IgG sensor chip for 200 s followed by 800 s of dissociation at a flow rate of 30 μL/min. The starting trimer concentration for each sample was 200 nM. Sensor chip surfaces were regenerated after each injection following manufacture instructions with Glycine 2.5 (GE Healthcare Life Sciences 10 mM glycine - HCl at pH 2.5) at a flow rate of 40 μL/min for 180 s.

### Mouse immunization

Mouse experiments were carried out in compliance with National Institutes of Health regulations and approval from the Animal Care and Use Committee of the Vaccine Research Center. Six week old female BALB/cJ mice (Jackson Laboratories) were inoculated intramuscularly with Sigma Adjuvant System, at weeks 0 and 3, as detailed previously^49^. Serum was collected 2 weeks post-prime and post-boost for measurements of antibody responses as detailed hereafter.

### Enzyme-linked immunosorbent assay (ELISA)

ELISA experiments were carried out as previously described^41^. Briefly, Nunc Maxisorp ELISA plates (ThermoFisher) were coated with 100 ng/well of stabilized soluble SARS-CoV-2 spike protein^24^ (with His-tag cleaved to remove potential cross-reactivity) in 1X PBS at 4 °C for 16 hr. To eliminate fold-on-specific binding, 50 μg/mL of fold-on protein was added to serial dilutions of heat-inactivated sera for 1 hr at room temperature (RT). After blocking in PBS-Tween (PBST) supplemented with 5% nonfat milk, plates were incubated with sera for 1 hr at RT. After blocking in PBS-Tween (PBST) supplemented with 5% nonfat milk, plates were incubated with serial dilutions of heat-inactivated sera for 1 hr at RT. Secondary antibody, goat anti-mouse IgG conjugated to horseradish peroxidase (ThermoFisher), was then added, followed by excitation with 3,5,3′5′-tetramethylbenzidine substrate (KPL). Each step in this procedure was followed by standard washes in PBST. Endpoint titers were calculated as the dilution factor that resulted in an optical density exceeding 4X background (secondary antibody alone).

### Lentivirus-based pseudovirus neutralization assay

The pseudovirus neutralization assay was performed as described previously^41,50^. To produce SARS-CoV-2 pseudovirus, a codon-optimized CMV/R-SARS-CoV-2 spike (Wuhan-1, Genbank #: MN908947.3) plasmid, was constructed and co-transfected with plasmids encoding luciferase reporter, human transmembrane protease serine 2 (TMPRSS2)^51^, and lentivirus backbone into HEK293T/17 cells (ATCC #CRL-11268), as previously described^52^. Heat-inactivated serum was mixed with the pseudovirus, incubated at 37 °C, and then added to ACE-2-expressing 293T cells. Cells were lysed after 72 hr, and luciferase activity was measured. Percent neutralization was calculated with uninfected cells as 100% neutralization and cells infected with only pseudovirus as 0% neutralization. ID_50_ titers were determined using a log (agonist) vs. normalized response (variable slope) nonlinear function in Prism v8 (GraphPad).

## Data availability

All relevant data are within the paper and its Supporting Information files.

## Acknowledgements

We thank T. Beaumont and H. Spits for antibody D25, A. Lanzavecchia for antibodies PIA174, PIA75 and MPE8, J. Stuckey for assistance with figures, and members of the Vaccine Research Center for discussions or comments on the manuscript. We thank members of the NIH NIAID VRC Translational Research Program for technical assistance with mouse experiments. Support for this work was provided by the Intramural Research Program of the Vaccine Research Center, National Institute of Allergy and Infectious Diseases, National Institutes of Health. This project has been funded in part with Federal funds from Frederick National Laboratory for Cancer Research, NIH, under Contract No. HHSN261200800001E (Y. Tsybovsky). K.S. Corbett is the recipient of a research fellowship that was partially funded by the Undergraduate Scholarship Program, Office of Intramural Training and Education, Office of the Director, NIH. The content of this publication does not necessarily reflect the views or policies of the Department of Health and Human Services, nor does mention of trade names, commercial products, or organizations imply endorsement by the U.S. Government.

## Author contributions

B.Z. and C.W.C. designed research with B.Z. heading protein design and production; B.Z., C.W.C., A.S.O., and R.V. produced nanoparticle and trimer proteins; C.W.C. and B.Z. prepared trimer-coupled nanoparticles and performed antigenic assessments; Y.T. performed negative-stain EM; S.W. assisted with manuscript assembly; T.Z. provided design for SARS-CoV-2 spike protein; G.S-J. provided the design for PIV3 F protein; A.P., L.W. and E.S.Y. provided pseudovirus; G.B.H. carried out immunizations; O.M.A. and A.W. performed ELISA; J.I.M. performed neutralization assay; B.S.G. and K.S.C. designed mouse experiment and oversaw ELISA experiment; J.R.M. oversaw pseudovirus preparation; N.J.S., B.S.G., and K.S.C. oversaw neutralization assay; E.P. and C.Y. performed data analyses for immunoassays; P.D.K. oversaw the project with B.Z., C.W.C., S.W., and P.D.K. writing the paper, and all other authors providing revisions and comments.

