## Supplementary material for "A Platform Incorporating Trimeric Antigens into Self-Assembling Nanoparticles Reveals SARS-CoV-2-Spike Nanoparticles to Elicit Substantially Higher Neutralizing Responses than Spike Alone": Table S1

**Supplementary Table S1. Amino acid sequences of constructs for protein expression.**

| Construct name | Amino acid sequence |
|---|---|
| LuS-N71-SpyTag | MDSKGSSQKGSRLLLLLVVSNLLLPQGVVGAHIVMVDAYKPTKGSGSAMQIYEGKLTAEGLRFGIVASRFNHALVDRL<br>VEGAIDAIVRHGGREEDITLVRVPGSWEIPVAAGELARKENISAVIAIGVLIRGATPHFDYIASEVSKGLADLSLELRKPIT<br>FGVITADTLEQAIERAGTKHGNKGWEAALSAIEMANLFKSLRGGLVPRGSHHHHHHSAWSHPQFEK |
| Ferritin-N96-SpyTag | MDSKGSSQKGSRLLLLLVVSNLLLPQGVVGQHHHHHHHHSAWSHPQFEKGGLVPRGGAHIVMVDAYKPTKGGGSG<br>DPMLSKDIIKLLNEQVNKEMQSSNLYMSMSSWCYTHSLDGAGLFLFDHAAEEYEHAKKLIIFLNENNVPVQLTSISAPE<br>HKFEGLTQIFQKAYEHEQNISESINNIVDHAIKSKDHATFNFLQWYVAEQHEEEVLFKDILDKIELIGNENHGLYLADQY<br>VKGIAKSRKS |
| RSV F-SpyCatcher | MELLILKANAITTILTAVTFCFASGQNITEEFYQSTCSAVSKGYLGALRTGWYTSVITIELSNIKEIKCNGTDAKVKLIKQE<br>LDKYKNAVTDLQLLMQSTPATGSGSAIASGVAVCKVLHLEGEVNKIKSALLSTNKAVVSLSGCGVSVLTFKVLDLKNYI<br>DKQLLPILNKQSCSIPNIETVIEFQQKNNRLLEITREFSVNAGVTTPVSTYMLTNSELLSLINDMPITNDQKKLMSNNVQI<br>VRQQSYSIMCIIKEEVLAYVVQLPLYGVIDTPCWKLHTSPLCTTNTKEGSNICLTRTDRGWYCDNAGSVSFFPQAETC<br>KVQSNRVFCDTMNSRTLPSEVNLCNVDIFNPKYDCKIMTSKTDVSSSVITSLGAIVSCYGKTKCTASNKCRGIIKTFSN<br>GCDYVSNKGVDTVSVGNTLYYVNKQEGQSLYVKGEPIINFYDPLVFPSDEFDASISQVNEKINQSLAFIRKSDELLSAIG<br>GYIPEAPRDGQAYVRKDGEWVLLSTFLGSGDSATHIKFSKRDEDGKELAGATMELRDSSGKTISTWISDGQVKDFYL<br>YPGKYTFVETAAPDGYEVATAITFTVNEQGQVTVNGKATKGDAHIGSGLVPRGSHHHHHHSAWSHPQFEK |
| PIV3 F-SpyCatcher | MYSMQLASCVTLTLVLLVNSQIDITKLQHVGVLVNSPKGMKISQNFETRYLILSLIPKIEDSNSCGDQQIKQYKRLLDRLII<br>PLYDGLKLQKDVIVTNQESNENTDPRTERFFGGVIGTIALGVATSAQITAAVALVEAKQAKSDIEKLKEAIRDTNKAVQS<br>VCSSVGNCIVAIKSVQDYVNKEIVPSIARLGCEAAGLQLGIALTQHYSELTNCFGDNIGSLQEKGIKLQCIASLYRTNITEI<br>FTTSTVDKYDIYDLLFTESIKVRVIDVDLNDYSITLQVRLPLLTRLLNTQIYKVDSISYNIQNREWYIPLPSHIMTKGAFLG<br>GADVKECIEAFSSYICPSDPGFVLNHEMESCLSGNISQCPRTTVTSDIVPRYAFVNGGVVANCITTTCTCNGIGNRINQ<br>PPDQGVKIITHKECNTIGINGMLFNTNKEGTLAFYTPDDITLNNSVALDPIDISIELNKVKSDLEESKEWYRRSNQKLSAI<br>EDKIEEILSKIYHIENEIARIKKLIGEAPGGSGGDSATHIKFSKRDEDGKELAGATMELRDSSGKTISTWISDGQVKDFYL<br>YPGKYTFVETAAPDGYEVATAITFTVNEQGQVTVNGKATKGDAHIGSGLVPRGSHHHHHHSAWSHPQFEK |
| SARS-CoV-2 spike-SpyCatcher* | MGWSCIILFLVATATGVHSAPELLGGPSVFLFPPKPKDTLMISRTPEVTCVVVDVSHEDPEVKFNWYVDGVEVHNAKT<br>KPREEQYNSTYRVVSVLTVLHQDWLNGKEYKCKVSNKALPAPIEKTISKAKGQPREPQVYTLPPSRDELTKNQVSLY<br>CLVKGFYPSDIAVEWESNGQPENNYKTTPPVLDSDGSFFLYSKLTVDKSRWQQGNVFSCSVMHEALHNHYTQKSLS<br>LSPGKGGGGSGGGGSGGGGSGGGGSAPELLGGPSVFLFPPKPKDTLMISRTPEVTCVVVDVSHEDPEVKFNWYVD<br>GVEVHNAKTKPREEQYNSTYRVVSVLTVLHQDWLNGKEYKCKVSNKALPAPIEKTISKAKGQPREPQVYTLPPSRDE<br>LTKNQVSLTCLVKGFYPSDIAVEWESNGQPENNYKTTPPVLDSDGSFFLTSKLTVDKSRWQQGNVFSCSVMHEALH<br>NHYTQKSLSLSPGKGGSGGGGSGGLEVLFQGPQCVNLTTRTQLPPAYTNSFTRGVYYPDKVFRSSVLHSTQDLFLP<br>FFSNVTWFHAIHVSGTNGTKRFDNPVLPFNDGVYFASTEKSNIIRGWIFGTTLDSKTQSLLIVNNATNVVIKVCEFQFC<br>NDPFLGVYYHKNNKSWMESEFRVYSSANNCTFEYVSQPFLMDLEGKQGNFKNLREFVFKNIDGYFKIYSKHTPINLV<br>RDLPQGFSALEPLVDLPIGINITRFQTLLALHRSYLTPGDSSSGWTAGAAAYYVGYLQPRTFLLKYNENGTITDAVDCA<br>LDPLSETKCTLKSFTVEKGIYQTSNFRVQPTESIVRFPNITNLCPFGEVFNATRFASVYAWNRKRISNCVADYSVLYNS<br>ASFSTFKCYGVSPTKLNDLCFTNVYADSFVIRGDEVRQIAPGQTGKIADYNYKLPDDFTGCVIAWNSNNLDSKVGGNY<br>NYLYRLFRKSNLKPFERDISTEIYQAGSTPCNGVEGFNCYFPLQSYGFQPTNGVGYQPYRVVVLSFELLHAPATVCGP<br>KKSTNLVKNKCVNFNFNGLTGTGVLTESNKKFLPFQQFGRDIADTTDAVRDPQTLEILDITPCSFGGVSVITPGTNTSN<br>QVAVLYQDVNCTEVPVAIHADQLTPTWRVYSTGSNVFQTRAGCLIGAEHVNNSYECDIPIGAGICASYQTQTNSPGSA<br>SSVASQSIIAYTMSLGAENSVAYSNNSIAIPTNFTISVTTEILPVSMTKTSVDCTMYICGDSTECSNLLLQYGSFCTQLNR<br>ALTGIAVEQDKNTQEVFAQVKQIYKTPPIKDFGGFNFSQILPDPSKPSKRSFIEDLLFNKVTLADAGFIKQYGDCLGDIA<br>ARDLICAQKFNGLTVLPPLLTDEMIAQYTSALLAGTITSGWTFGAGAALQIPFAMQMAYRFNGIGVTQNVLYENQKLIA<br>NQFNSAIGKIQDSLSSTASALGKLQDVVNQNAQALNTLVKQLSSNFGAISSVLNDILSRLDPPEAEVQIDRLITGRLQSL<br>QTYVTQQLIRAAEIRASANLAATKMSECVLGQSKRVDFCGKGYHLMSFPQSAPHGVVFLHVTYVPAQEKNFTTAPAIC<br>HDGKAHFPREGVFVSNGTHWFVTQRNFYEPQIITTDNTFVSGNCDVVIGIVNNTVYDPLQPELDSFKEELDKYFKNHT<br>SPDVDLGDISGINASVVNIQKEIDRLNEVAKNLNESLIDLQELGKYEQGSGYIPEAPRDGQAYVRKDGEWVLLSTFLGR<br>SGGGLVPQQSGDSATHIKFSKRDEDGKELAGATMELRDSSGKTISTWISDGQVKDFYLYPGKYTFVETAAPDGYEVA<br>TAITFTVNEQGQVTVNGKATKGDAHI |

* This amino acid sequence includes a single chain Fc purification tag (see reference 38).
